# Insights on Human Dimensions of Freshwater Fish Conservation in Jharkhand and Bihar, India

**DOI:** 10.64898/2026.02.10.705004

**Authors:** Prantik Das, V. V. Binoy

## Abstract

Conservation outcomes in the socio-economically disadvantaged regions are strongly influenced by human behaviour, social norms, and existing governance mechanisms. This study examined stakeholder attitudes, perceptions, values, norms and decision-making processes associated with the conservation of freshwater fishes in two neighbouring states in Eastern India - Jharkhand and Bihar. An approach integrating the Conservation Planning Framework (CPF) with the Theory of Planned Behaviour (TPB) and Social Values (SV) enabled the development of four interlinked themes: “livelihood and economic prioritisation over conservation”, “constraints on participation”, “values and conservation willingness” and “erosion of social and cultural memories of mahseer”, indicative of a process of dual extinction faced by these iconic freshwater fishes. Despite the widespread positive attitudes of the stakeholders towards native fishes and freshwater ecosystems, conservation intentions and actions in both states were found to be negatively influenced by feeble communication, prioritisation of aquaculture, institutional rigidity, inadequate conservation education, limited actual behavioural control (ABC) and subjective norms-driven livelihood pressures. However, the presence of active fishermen cooperative societies and stronger relational values among the local communities makes Jharkhand better equipped to implement participatory governance and stakeholder-involved conservation engagement plans. By strategically linking CPF, TPB, and SV, this study demonstrates how human attitudes, behaviour, social norms, and institutional structures interact to shape freshwater fish conservation outcomes in regions where livelihood needs intersect with conservation priorities, thereby offering actionable insights for managing the native freshwater fish diversity.

## 1. Introduction

Freshwater ecosystems, lifelines for both biodiversity and human societies, are among the most imperilled and threatened habitats on earth (He et al. 2017; Reid et al. 2019; Arora et al. 2024; Raghavan et al. 2025). Despite occupying a relatively small fraction of the earth‘s surface, they support a disproportionately high share of global biodiversity and underpin food security and livelihoods for millions. Amongst the countless species surviving in these water-bodies, fishes are particularly vulnerable. Freshwater fishes are recognised as the most threatened vertebrate group, with extinction rates estimated to be five times higher than those of terrestrial fauna and three times higher than those of marine mammals (Duncan and Lockwood 2001; Dias et al. 2017; Vardakas et al. 2025). Decline in the freshwater fish diversity is accelerated by multiple interacting pressures - climate change and human induced habitat degradation and fragmentation, river regulation and flow alteration by dams and barrages, water pollution, overexploitation, and introduction of invasive alien fish species (IAFs; Lakra et al. 2010; Dudgeon 2011; Reid et al. 2019).

In the densely populated river basins of the Global South, both ecosystems and fishes are subject to sustained and intensive human exploitation (Dudgeon 2011; Kelkar 2014; Barletta et al. 2015). The rapid and extensive transformations happening in the rivers and its watershed over the last many decades (Welcomme et al. 2010; Lynch et al. 2016) has led to the degradation of the quality of these aquatic resources and the hence the strength and diversity of fish populations. The limited availability or absence of long-term records of fish catches, fishing intensity, gear types used, and population trends of the exploited species (Arthington et al. 2004; Bartley et al. 2015) and the need for considering numerous human factors manifolds the complexity of the conservation efforts in these socio-ecosystems. The Gangetic River basin in India, one of the world‘s largest and most heavily utilised river systems (Kelkar 2014; Tandon and Sinha 2018), supports remarkable freshwater biodiversity and offers many essential natural resources and livelihood for millions (Sarkar et al. 2012; Kelkar 2014; Bharati et al. 2016). The eastern region of the Gangetic basin, encompassing the Indian state of Bihar and parts of neighbouring Jharkhand, is of particular ecological and economic significance due to its dense network of rivers and tributaries such as the Ganga, Kosi, Sone, Bagmati, Gandak, Burhi Gandak, Damodar, Barakar, Ajay, Punpun, North Koel, etc. (Kelkar 2014; Dey et al. 2020; Kumari and Sharma 2022; Kumar, V. et al. 2025). Additionally, several other rivers originating in Jharkhand, such as the Kharkai, Subarnarekha, and South Koel, flow eastward and drain into separate basins, namely the Subarnarekha and Brahmani river basins, respectively (Tudu and Maji 2025).

The floodplains, wetlands, streams, hill and upland rivers, and reservoirs constitute a dynamic freshwater landscape that historically supported high levels of native fish diversity (Kumar, V. et al. 2025). Several commercially important piscine species such as rohu (*Labeo rohita*), catla (*Catla catla*), mrigal (*Cirrhinus mrigala*), magur (*Clarias magur*; State Fish of Jharkhand), stinging catfish (*Heteropneustes fossilis*; State Fish of Bihar) support local communities through riverine capture and inland fisheries contributing to livelihoods, income, employment, and food security (Kumar et al. 2019; Kumari and Sharma 2022; Kumar, V. et al. 2025; Tudu and Maji 2025. The presence of the mahseer (*Tor* spp. and *Neolissochilus* spp.) iconic freshwater fish with ecological, socio-cultural, religious and economic importance (Bhatt and Pandit 2016; Pinder et al. 2019; Sarma et al. 2022; Froese and Pauly 2025) heightens the conservation importance of this region. Archival literature, gazetteers, research articles, and historical records document the presence of four species of mahseer in these two states, viz. *T. putitora* (Himalayan or golden mahseer; Endangered), *T. tor* (red-fin or deep-bodied mahseer; Data Deficient), *T. mosal* (mosal mahseer; Data Deficient), and *N. hexagonolepis* (chocolate mahseer; Near Threatened) (O‘Malley 1906, 1910, 1924, 1926; Roy Choudhury 1957; Kumar 1970; Das 2005; Srivastava and Singh 2014; Sinha Ray 2016; Tudu and Maji 2025). Additionally, the Kosi River in Bihar is recognised as the type locality of *T mosal* (Raghavan et al. 2017), a species for which no recent voucher specimens are available from the region (Dahanukar et al. 2018; Pinder et al. 2019).

Despite their high conservation value, increasing habitat degradation and fragmentation, freshwater ecosystems of Bihar and Jharkhand and the fish diversity present there received relatively little conservation attention (Srivastava and Singh 2014; Kelkar 2014; Das et al. 2018; Arora et al. 2024; Pandey and Bharti 2024). Over the recent decades, embankment construction (*bandh*), river regulation, sand, boulder and river-bed mining, pollution, destructive fishing practices such as the illegal gear use, netting, dynamiting, electrocution, poisoning, and the spread of invasive alien fish species (IAFs) has ended up in habitat simplification, loss of spawning grounds, disruption of migratory pathways, etc. igniting widespread declines in the native riverine fish populations in these states (Singh and Lakra 2006; Sarkar et al. 2012; Kelkar 2014; Goswami 2023; Das and Khan 2024; Pandey and Bharati 2024; Prasad 2025). The local fishing communities depend on subsistent fishing of species such as climbing perch (*Anabas testudineus*), snakeheads (*Channa* spp.), and bronze featherback (*Notopterus notopterus*), Indian featherback/Indian knifefish (*Chitala chitala*), minor carps and barbs (*Puntius* spp.), eels such as Indian Mottled Eel (*Anguilla bengalensis*), minnows, and Sucker head fish (*Garra gotyla*) for livelihood are also suffering from decline in fish populations. Although ecological surveys, distributional analyses, and taxonomic studies have generated valuable insights into fish diversity and habitat degradation in several areas of Bihar and Jharkhand (Kumar, V. et al. 2025; Tudu and Maji 2025) very little attention has been given to the human dimensions shaping people‘s interaction with freshwater fishes and ecosystems, and their responses to conservation initiatives in these states.

It is well known, in human-dominated ecosystems including freshwaters, conservation outcomes are strongly influenced by the attitudes, perceptions, values, norms, and behaviours of numerous stakeholders involved (Johannes et al. 2000; Silvano and Valbo-Jørgensen 2008; Bennett et al. 2017; Pascual et al. 2023). While some studies from Bihar have highlighted the value of incorporating fishermen‘s knowledge in understanding fish declines and environmental changes (Dey et al. 2020), its translation into conservation action requires data of the mindset of a spectrum of actors, nature of power relations existing between them, subjective norms, values, economic drivers and institutional constraints prevailing in the society. This imbalance present in the essential information serves as a major barrier for designing conservation strategies that are ecologically effective, socially acceptable and livelihood-linked (Kareiva and Marvier 2012; Marchini 2014; Dacks et al. 2025). Addressing this gap the present study examined different facets of freshwater fish conservation in Jharkhand and Bihar through a socio-behavioural lens focusing on the stakeholders‘ perceptions, attitudes, values, and behaviours related to freshwater ecosystems and fish conservation. In order to get an illustrated picture, we followed an approach integrating Conservation Planning Framework (CPF; Marchini et al. 2019), the Theory of Planned Behaviour (TPB; Ajzen 1985; 1991), and Social Value systems (SV; Chan et al. 2016; Pascual et al. 2023; Das and Binoy 2025b). Here, the CPF provides a well-structured, system-wide approach for assessing the present conservation context, identifying problems, setting objectives, and evaluating management options within the complex socio-ecological systems. The second element TPB, elucidates how attitudes, subjective norms, and perceived and actual behavioural control shape conservation-relevant behaviours (Ajzen 1985; 1991), while the Social Value (SV) captures the intrinsic (valuing wildlife and nature for their own sake and inherent worth, independent of human use or benefit), instrumental (valuing wildlife and nature based on human material needs and usefulness; valuing them for tangible benefits like livelihoods), and relational values (values for wildlife developed through cultivated human-nature relationships and meanings, encompassing a sense of morality, spirituality, identity, belonging, and personal as well as cultural connections with wildlife) underpinning human-fish relationships, including moral, stewardship and responsibility-based aspects for effective human-animal coexistence (Chan et al. 2016; Pascual et al. 2023). Such an integrated approach has been successfully utilised to get insights into the conservation of freshwater fishes mahseers in the states of Karnataka, Assam and Uttarakhand very recently (Das and Binoy 2025b) In this scenario by keeping the selected districts of the Indian states of Jharkhand and Bihar in focus, the present study aims to:

1. Understand the attitudes, values, perceptions of various stakeholders towards freshwater fish species, their conservation and ecosystem management in Jharkhand and Bihar using the integrated CPF-TPB-SV framework.
2. Explore the implications of these findings to develop recommendations for context-specific, socially informed freshwater fish conservation strategies for these states.

## 2. Methodology

### 2.1. Study Areas

This study was conducted across two adjoining states in Eastern India, Jharkhand and Bihar. Bihar encompasses extensive lower reaches of the River Ganga, as well as a major portion of the River Kosi, a key tributary within the Gangetic River Basin. Additionally, the Eastern and Northern regions of Jharkhand fall within the Gangetic River Basin, while other catchments originating in the Chota Nagpur Plateau region of Jharkhand drain into multiple river systems, including the Gangetic, Subarnarekha, and Brahmani River Basins (Jain et al. 2007; Singh and Giri 2018; Tudu and Maji 2025). The fieldwork was undertaken in eight districts of Jharkhand viz., Dhanbad, Dumka, Gumla, Hazaribagh, Koderma, Latehar, Lohardaga and Ranchi (Fig. 1), and in three districts of Bihar viz., Patna, Saharsa and Supaul (Fig. 2). These districts were selected to capture the geographical, ecological, hydrological, socio-cultural, and economic heterogeneity present across the two states. Major rivers flowing through the districts studied were Son, North Koel, South Koel, Sankh, Damodar, Barakar, Subarnarekha, Mayurakshi, and Ajay in Jharkhand and Ganga, Kosi, Son, and Punpun in Bihar.

**Fig. 1.**
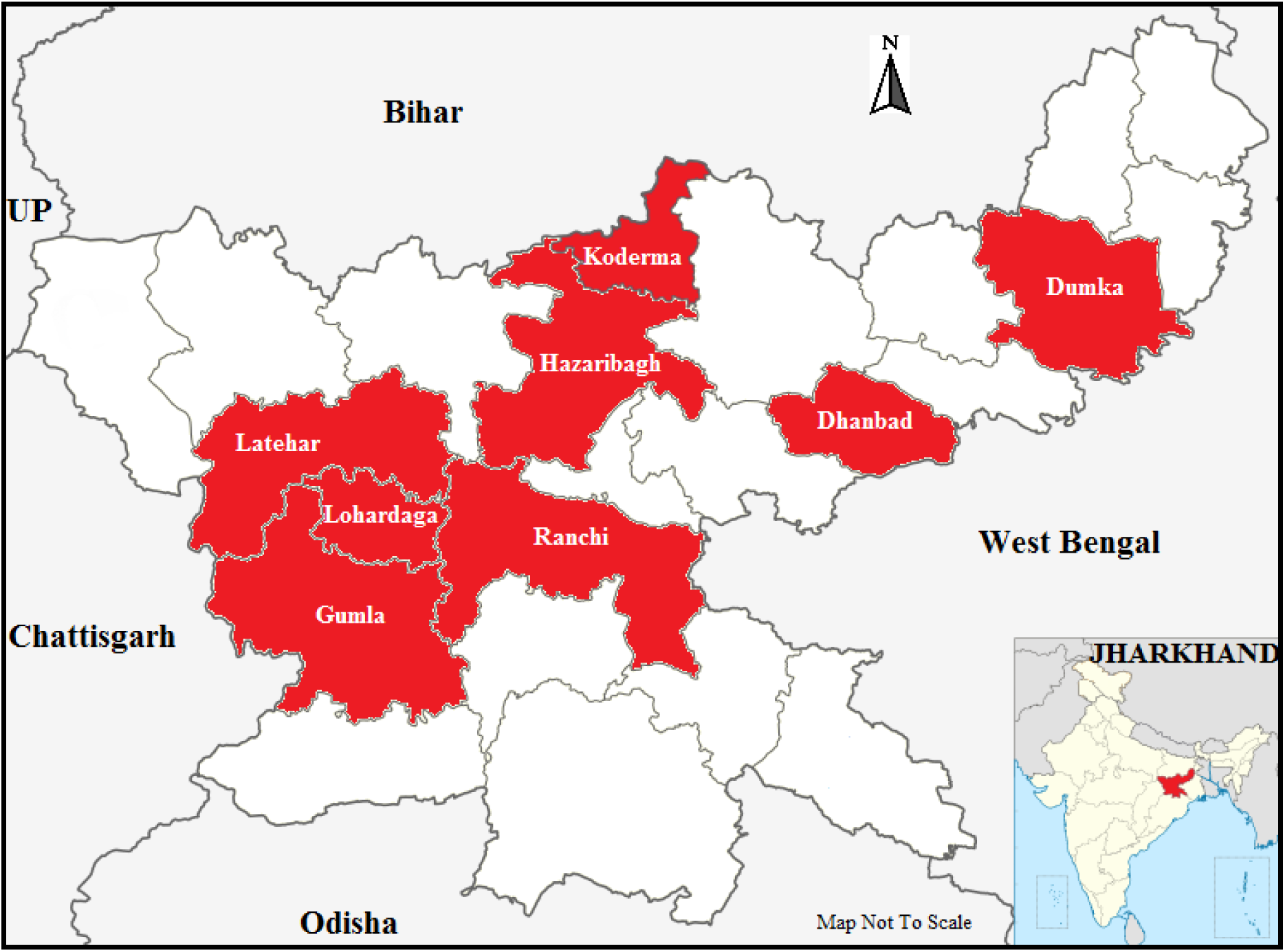
The districts in the Indian state of Jharkhand where the study was carried out. Map not to scale.

**Fig. 2.**
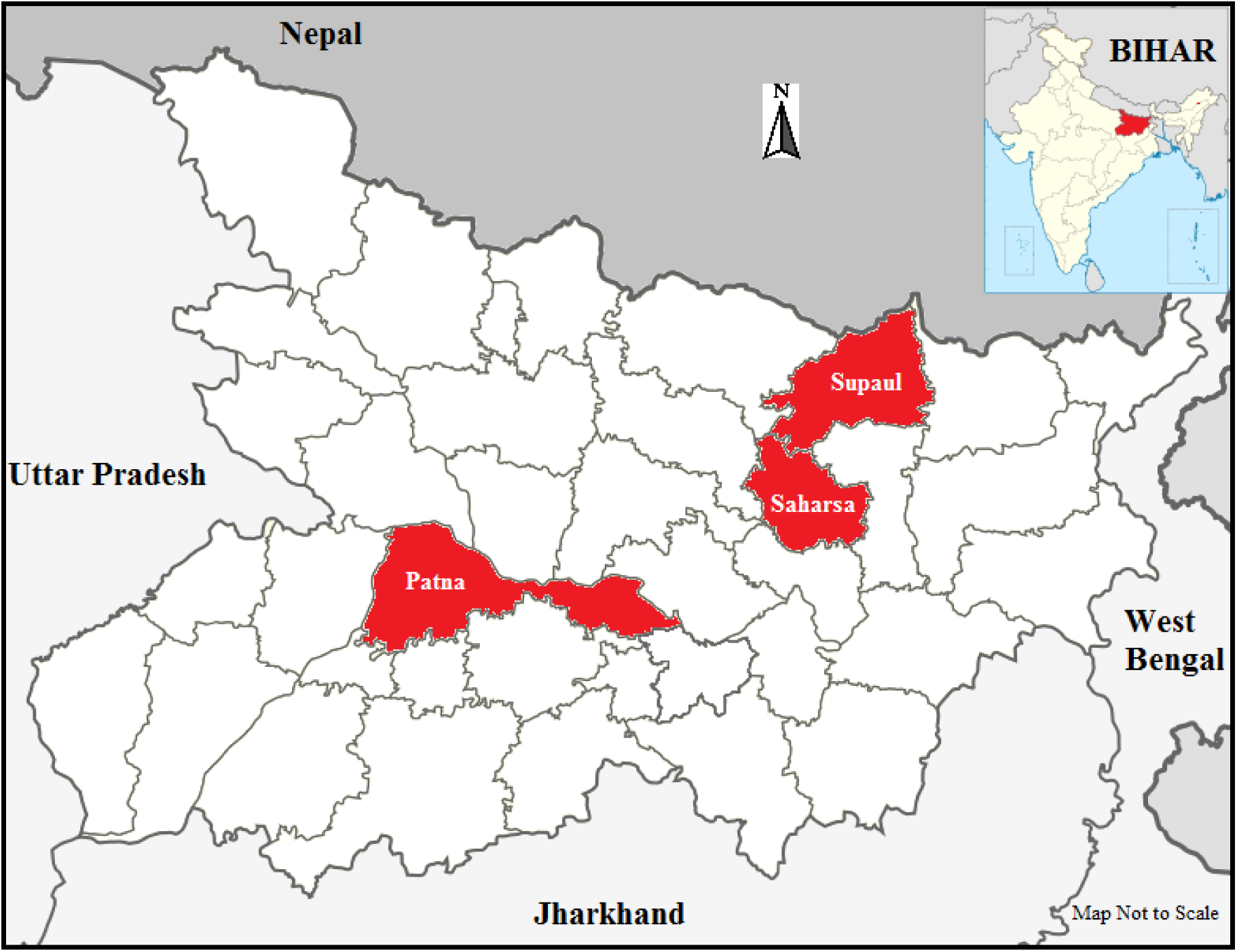
The districts in the Indian state of Bihar where the study was carried out. Map not to scale.

### 2.2. Data Collection

Key stakeholder groups were identified using purposive sampling, followed by snowball sampling to reach additional resource persons with relevant experience and knowledge of freshwater fishes and ecosystems. In-depth, semi-structured, discursive, one-on-one interviews of the stakeholders were conducted following a list of questions reflecting multiple aspects of the integrated CPF-TPB-SV framework (SM 1), which lasted between 15-30 minutes. Perceptions and attitudes towards freshwater fishes and riverine ecosystems, environmental changes observed in such habitats, attitudes and values toward conservation, conservation communication, environmental decision-making strategies, determinants of the behavioural control, subjective and social norms, and barriers of pro-conservation and pro-environmental actions were the topics covered during the stakeholder interaction. In total, 69 and 84 interviews were conducted in Jharkhand and Bihar respectively (Table 1). In Jharkhand the languages used for the communication were Khortha, Hindi and Bangla, while stakeholders were comfortable with Hindi, Urdu and English in Bihar Interviews were audio recorded, written notes taken and photographs captured after acquiring oral/audio-recorded informed consent from the participants. The participants were assured of the anonymity of their names, personal details and designations to protect their identity and privacy. We ensured that all interviews adhered to the required ethical guidelines for conducting non-invasive human research.

**Table 1.**
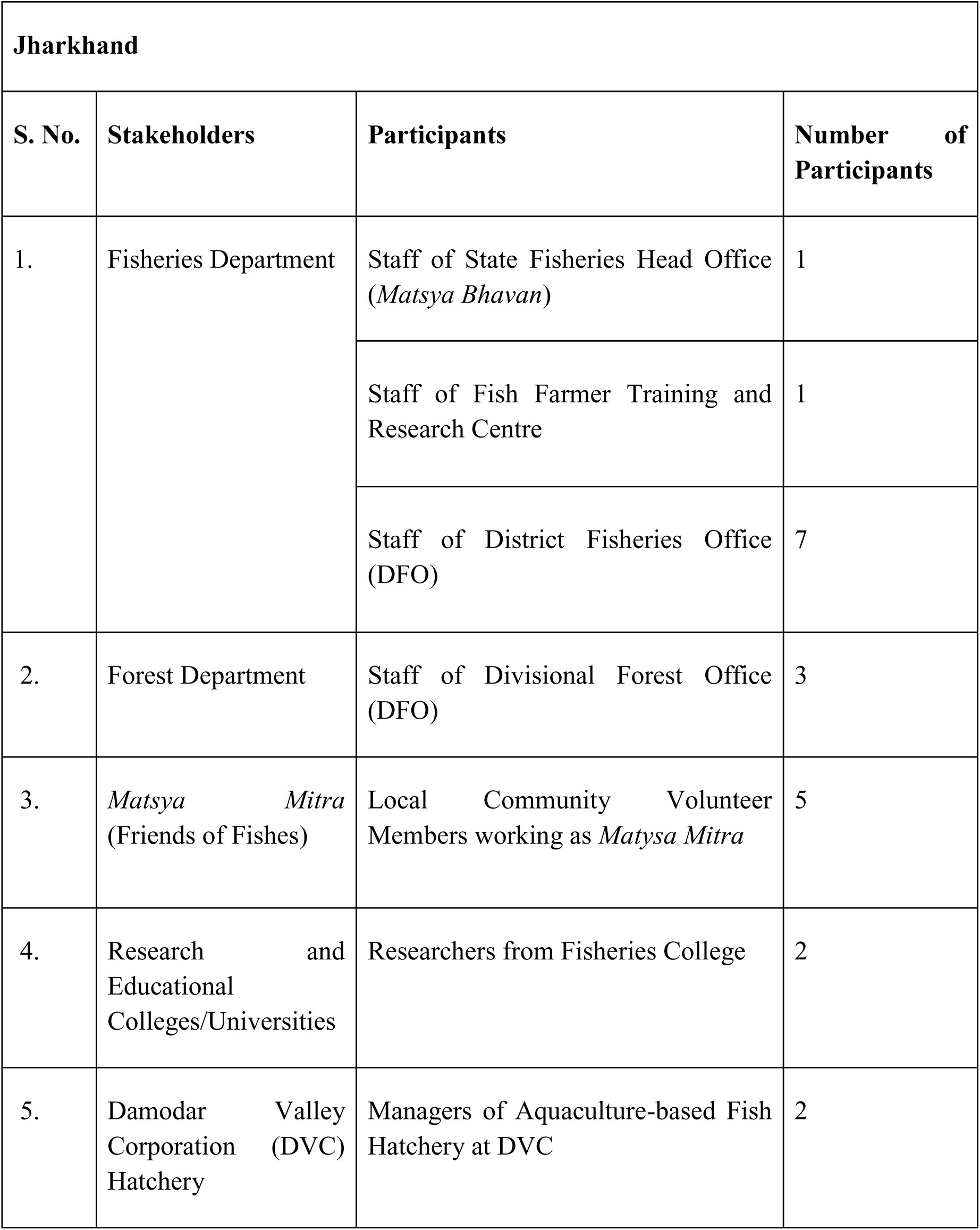
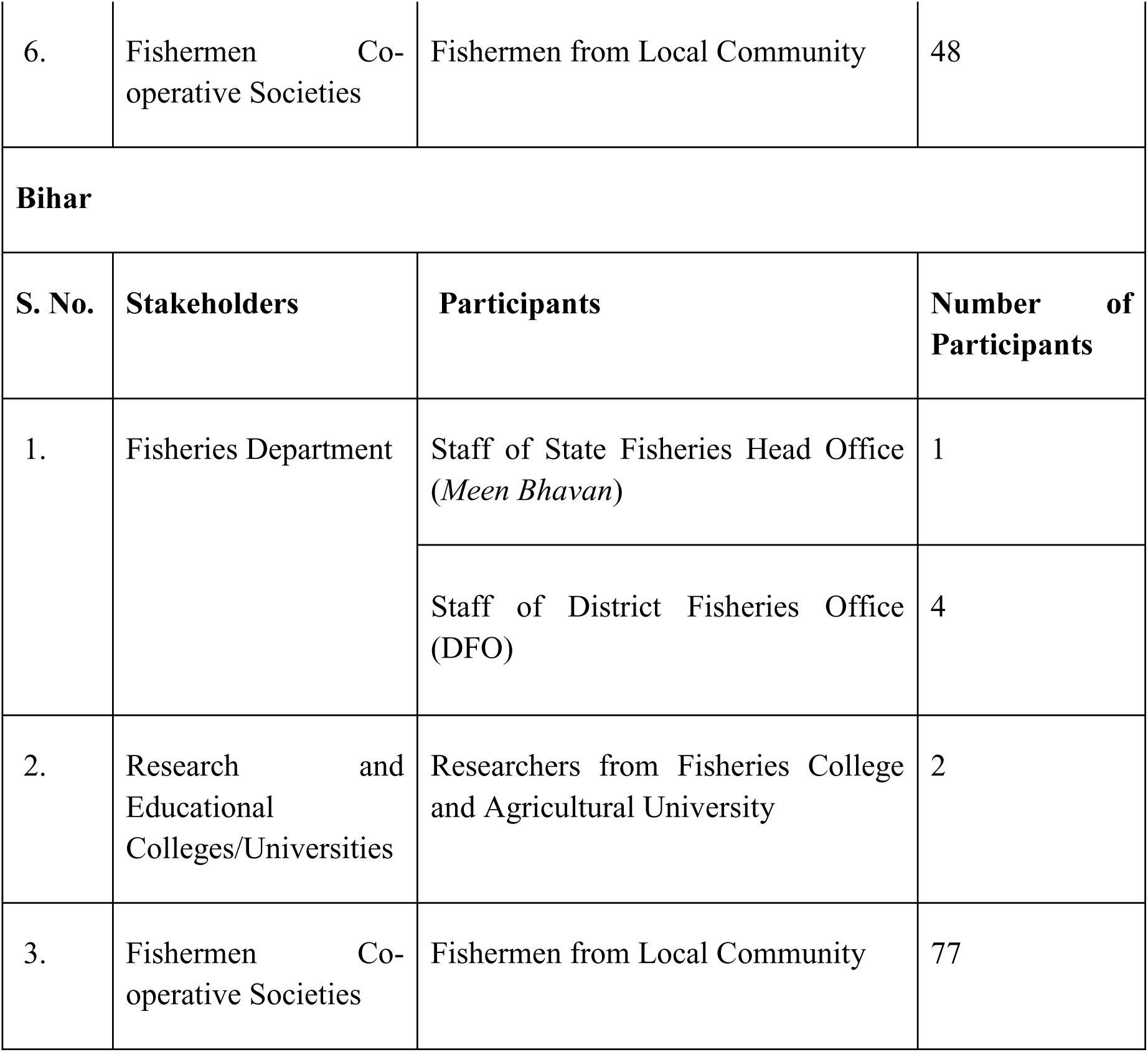
List of stakeholders from the two Indian states interviewed for the study.

### 2.3. Data Analysis

The interviews conducted in local languages were transcribed verbatim into English through manual transcription, while interviews conducted in English were transcribed using Rev.ai. All transcripts were subsequently cross-checked manually against the original audio recordings and supplemented with field notes to correct inconsistencies in transcription and to retain contextual nuance (Nkansah-Dwamena 2023). The qualitative data collected were analysed using Reflexive Thematic Analysis (RTA) with a hybrid deductive-inductive coding strategy to enhance analytical depth and rigour (Fereday and Muir-Cochrane 2006; Braun and Clarke 2013, 2019, 2021; Braun et al. 2014, 2022; Byrne 2022; Proudfoot et al. 2022; Das and Binoy 2025b). Following Braun and Clarke‘s (2006) six-step framework, the RTA involved data familiarisation, initial code generation (combining deductive and inductive codes), organisation of codes into meaningful themes, review and refinement of themes, and final definition of themes. Deductive codes were guided by the constructs from the integrated conceptual framework, whereas inductive codes captured the additional, context- and stakeholder-specific insights. This reflexive approach centres the researcher‘s subjectivity, knowledge, and contextual expertise and thoughtful qualitative interpretation. Accordingly, RTA, unlike the more conventional qualitative data coding approaches, does not rely on inter-coder reliability or replicability as measures of validity (Braun and Clarke 2019, 2021). Consequently, the entire dataset was coded by a single researcher (PD) with extensive familiarity with the methods of the RTA and the topic of enquiry.

## 3. Results

The first theme developed at the juncture of CPF‘s situation assessment with the components of TPB (attitude, subjective norms, PBC and ABC; Das and Binoy 2025b), was “livelihood and economic prioritisation over conservation.” Across both states, the majority of the stakeholders viewed the fishes primarily through a lens of livelihood (Jharkhand - J: 69; Bihar - B: 82). Words of a member of the local fishermen cooperative society in Jharkhand capture this situation precisely.

> Quote 1: “We mostly breed and farm IMCs and other fishes like common carp, grass carp, silver carp. We grow them in our ponds and lakes. Once they are big enough, we sell them. This is our livelihood. Yes, all native fishes should be conserved. We are tribal people, who worship nature and consider fishes to be a part of nature, but who has so much time to invest in such activities completely? We have to earn; we have to feed our families. We make a living by selling the fishes we grow.”

Although respondents were aware of the negative impact of the IAF on native species (J: 69, Fig. 3; B: 84, Fig. 4), many still considered these fishes (e.g. common carp, Thai magur, and tilapia, etc.; Canonico et al. 2005; Singh and Lakra 2006), along with other aquaculture species such as Indian Major Carps (IMC) to be their economic backbone (J: 64; B: 82). While Jharkhand banned wild stocking of the IAF tilapia in 2012–2013, controlled cage culture (16) of mono-sex Genetically Improved Farmed Tilapia (GIFT) strains with apt biosecurity measures (Press Information Bureau, Government of India 2024) is currently allowed in the state.

**Fig. 3.**
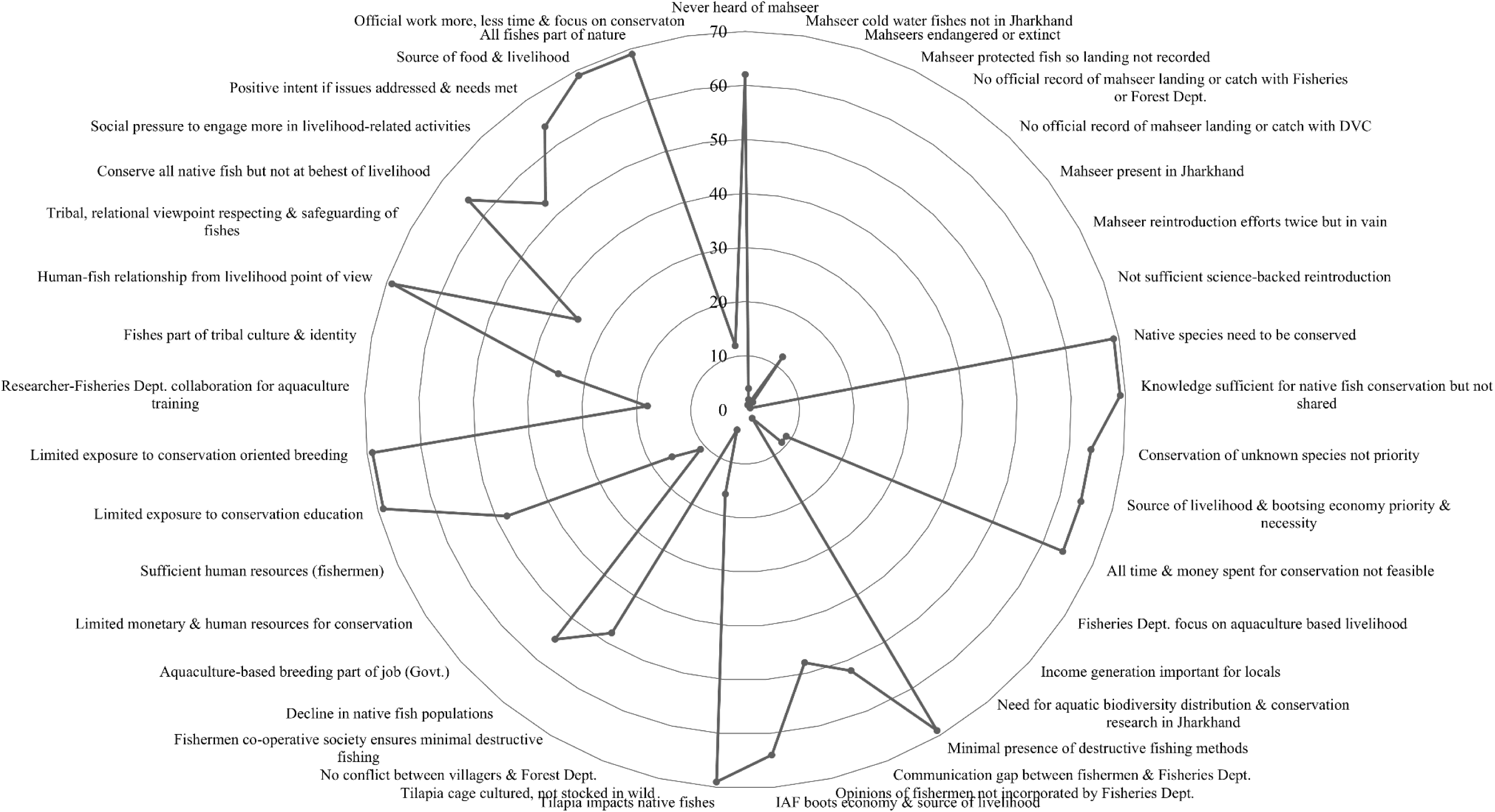
Radar chart depicting the frequency of codes obtained from the reflexive thematic analysis conducted on the interviews from Jharkhand.

**Fig. 4.**
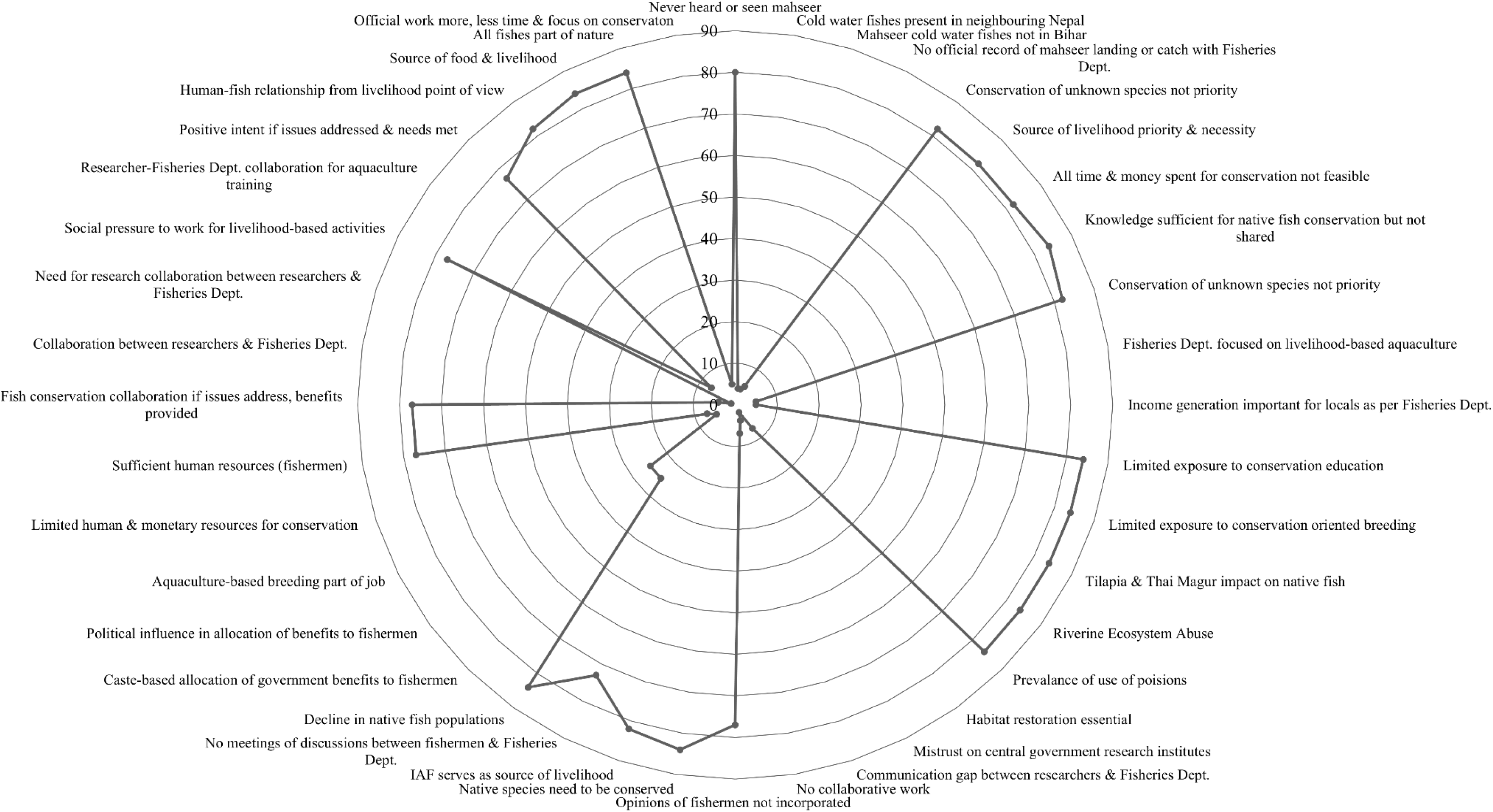
Radar chart depicting the frequency of codes obtained from the reflexive thematic analysis conducted on the interviews from Bihar.

For the fisheries departments of both states also conservation remained second to aquaculture promotion and livelihood enhancement (J: 9; B: 5).

> Quote 2: “We primarily promote aquaculture and farming of Indian Major Carps, as well as other fishes like pangasius, common carp, grass carp, etc., and encourage local people to take up fish cultivation. Over the years, fish production has increased, and fish farming has become one of the most important income-generating activities in Jharkhand, significantly reducing poverty levels. It is helping many local people earn a livelihood and is also boosting the state’s economy.” – said one of the fisheries department staff from Jharkhand.

Poor economic conditions and the vulnerability of people dependent on fishes for their income, visible across both states, may be contributing to the popularisation of strong subjective norms (a component of TPB), creating social pressure at the individual level to prioritise livelihood security over fish conservation (J: 53; B: 77). Such conditions, which reinforce attitudes supporting a “production-over-protection” approach, can constrain stakeholders‘ behavioural intentions. Perceived inadequate institutional and governmental support for conservation, along with the absence of alternative livelihood opportunities for those engaged primarily in conservation activities, is reflected in limited actual behavioural control (ABC-TPB). Consequently, involvement in conservation activities are often viewed as economically risky and a luxury rather than a necessity by many respondents (J: 64; B: 82). This perception was further compounded by the limited exposure to conservation-focused education and training in both states (J: 69; B: 84), where capacity building is centred around aquaculture, breeding techniques and production-enhancing technologies, possibly promoting a more utilitarian outlook towards freshwater fish species.

The decision-making and implementation elements of CPF with the attitude and PBC of TPB led to the next theme: “constraints on participation”. A top-down decision-making and governance structure with minimal possibility for stakeholder participation and communication between them hindered inclusive fish conservation in the focal states. We noticed significant communication gaps existing between fishermen, and fisheries department staff (J: 52; B: 73) and researchers and fisheries department (B: 4). Many fishermen from both states complained of their exclusion from decision-making processes (B: 77; Fig. 4) and neglect of their suggestions (J: 48; Fig. 3) by the authorities. The words of the member of a local fishermen‘s cooperative society in Bihar vouch for it.

> Quote 3: “We have requested meetings multiple times. I have even personally sent letters to the Fisheries head office, offering our help to address the issue of destructive illegal fishing. But to this day, there has been no response from them.”

However, instances of limited collaboration and joint activities, though restricted to aquaculture-related training and IMC breeding (J: 18; B: 7), particularly between researchers and the fisheries department staff were observed in both states. Unfortunately, many respondents who participated in such initiative were dissatisfied with the lack of interest for continued collaboration and bidirectional communication shown by the participants from other departments, as exhibited in the statement by a member of the fisheries department staff in Bihar:

> Quote 4: “I have told the senior officials many times that researchers from central institutes should consult us, the local government officials, on how to undertake breeding ground revival and habitat restoration, and how to protect rivers and their biodiversity by removing the bandhs/bunds. We know these things better; we have both the ground-level experience as well as knowledge from the fisheries research books we have read. But who cares? They just hear us as an obligation and never implement anything we suggest. So, we have stopped providing recommendations anymore.”

Such a situation can catalyse distrust between the departments (2), suppress conversation intentions among stakeholders, further weaken collaborations (7) and ultimately impact conservation actions as observed in Bihar.

Reduced availability of human and monetary resources also constrained actual behavioural control (ABC) of many stakeholders in both Jharkhand (16) and Bihar (7). For instance, many government staff expressed anguish over lesser time available to focus on conservation due to their heavy workload (J: 12; B: 5). Formally involving fishermen who hold pro-conservation and relational values for native fishes in the conservation intervention could fill this gap, and such communities acknowledged availability of the suitable human resource with them for this purpose in both Jharkhand (48) and Bihar (77). Unfortunately members from fishermen communities failed to join such activities due to lack of conservation education (J: 69; B: 84), limited exposure to conservation breeding initiatives (J: 69; B: 84), and exclusion from institutional decision-making and planning processes. In Bihar, caste bias and political influences affected fishermen‘s access to the government schemes (*yojanas*) and benefits (25). This inequity in the resource distribution was found emerging as a major barrier for participation in the fish conservation activities by many stakeholders. Hence, strengthening government support mechanisms, fulfilling needs, and addressing livelihood concerns (PBC; J: 64; B: 77) of the local populations may improve their willingness to join such activities. The following words of a member of the local fishermen cooperative society in Bihar reflected the same.

> Quote 5: “If proper benefits from the government schemes like fish feed, medicines, tube well, pump, aerators equipment, subsidies on pond construction and infrastructure, etc. are provided to us; then of course, why would we not help them in whatever conservation activity they ask us to do. All fishes should be conserved, but if we are unable to earn and don’t receive the benefits from the schemes, why should we invest our time in something that does not feed our families? They should stop this caste- and politics-based distribution and provide equal benefits to everyone.”

Insights from the monitoring and evaluation stages of CPF, combined with stakeholder‘s perceived and actual control of behaviours and subjective norms (TPB), and the 3 components of social values (SV) resulted in the theme “values and conservation willingness.” Across both states, all stakeholders consistently expressed strong intrinsic and relational values for fishes, underpinned by the belief that “*jungli*‘ (native wild species) are integral components of nature (J: 69; Fig. 3; B: 84; Fig. 4) requiring protection (J: 69; B: 84). Interestingly these perspectives coexisted with instrumental value - viewing fishes also as a source of food, livelihood, and income (J: 69; B: 84) - reflecting a pluralistic value orientation ascribed by the stakeholders toward native fishes. In Jharkhand, fishermen disclosed elements of the local and tribal identity, culture, and traditional belief systems (35) emphasising respect for and promoting safeguarding native fishes (35). Consistent with these value orientations, a much lower number of reports of destructive fishing received from this state (69). Here, the fishermen‘s cooperative societies, possibly reinforced by the stewardship virtues and relational outlook towards fish and nature (48) monitored and protected the aquatic resources, as indicated by one of their members.

> Quote 6: “Here, destructive fishing practices in the rivers and streams are very less. We do hear about such things happening in other states, but we are traditional fishermen, and as a cooperative society, we feel that it is our duty and responsibility to look after and protect our water bodies. Our livelihoods have come from these waters for generations. All fishes are part of nature; and nature has been providing us with food for generations. That is why we actively make sure that no one puts dynamite, poisons or chemicals into the rivers. Yes, there have been one or two small instances of fine-mesh or prohibited netting that happens sometimes, but otherwise, such cases are rare here.”

By contrast, despite the fishermen in Bihar expressing strong relational values towards native fishes (77), destructive fishing practices were widely reported (84). These practices included poisoning, use of chemicals, fertilizers, fine-mesh nets, dumping of sewage and solid waste along approximately 35-40 km stretches of the Kosi River, (84) which were allegedly carried out by the local non-traditional fishermen. Additionally, in both states staff of the government department (fisheries department in both states and forest department in Jharkhand) had an ‗instrumental‘ outlook towards fishes (J: 11; B: 5) and many described aquaculture as their prior duty.

Despite widespread relational values and pro-conservation orientations, in both states stakeholder‘s reported familiarity with native fishes was largely restricted to aquaculture- and livelihood-relevant species (Table 2). Interestingly, ecologically important but economically less-significant native species (Ayyappan and Raman 2003; Bhakta 2020; Baitha et al. 2024; Sahil et al. 2024), such as mustached danio (*Danio dangila*), finescale razorbelly minnow (*Salmostoma phulo*), large razorbelly minnow (*Salmostoma bacaila*) Indian flying barb (*Esomus danricus*), sucker-head fish (*Garra gotyla*), cotio (*Osteobrama cotio*), etc. were rarely recognised or reported by most of the participants, except managers of Damodar Valley Corporation fish hatchery in Jharkhand (Table 2). This knowledge gap was at its peak in the case of a socio-culturally significant but threatened group of fishes, the mahseer (*Tor* and *Neolissochilus spp*.). Across both states, lion‘s share of stakeholder groups interviewed had never heard of mahseer or were familiar with their vernacular names - *mahasol, turiya, mansir, masal* and *kajur* and failed to identify the fish from the images shown (J: 62; Fig. 3; B: 80; Fig. 4). However, mahseers were widely distributed in the freshwaters of the two focal states as mentioned and documented in the British Gazetteers during the colonial period and Bihar and Bengal Gazetteers post-independence (O‘Malley 1906, 1910, 1924, 1926; Roy Choudhury 1957; Kumar 1970; Talwar and Jhingran 1991; Sugnnan 1995; Desai 2003; Das 2005; Srivastava and Singh 2014; Sinha Ray 2016). This loss of collective knowledge and lived experience associated with mahseer led to the development of another theme - “erosion of social and cultural memories”. The message conveyed in the following statement by a member of the fishermen co-operative society in Bihar, was a representation of the stakeholder response regarding mahseers across Bihar and Jharkhand.

> Quote 7: “I have never heard of mahseer fish. (After looking at the picture) No sir, this type of fish is not found in the rivers here. I have never seen or caught any fish like this”.

**Table 2.** Vernacular and scientific names of the freshwater fish species reported by different stakeholder groups as occurring or present in the water bodies of Jharkhand and Bihar.

| S. No. | Vernacular Name (s) | Scientific Name | Stakeholder Groups Reporting the Species |
| --- | --- | --- | --- |
| 1. | Koi (Jharkhand)<br>Kowai, Kawai, Kabai (Bihar) | *Anabas testudineus* | All |
| 2. | Baamacchh | *Anguilla bengalensis* | Staff of the Fisheries Department, Bihar<br>Members of the Fishermen Co-operative Society, Bihar |
| 3. | Baghair, Baghar | *Bagarius* spp. | All |
| 4. | Katla | *Catla catla* | All |
| 5. | Garai | *Channa punctatus* | All |
| 6. | Shol, Saur | *Channa striatus* | All |
| 7. | Chital, Bhuna | *Chitala chitala* | All |
| 8. | Mrigal | *Cirrhinus cirrhosus* | All |
| 9. | Magur | *Clarias magur* | All |
| 10. | Bachwa, Garua | *Clupisoma garua* | All |
| 11. | Grass carp | *Ctenopharyngodon idella* | All |
| 12. | Pahari (Jharkhand)<br>Common carp | *Cyprinus carpio* | All |
| 13. | Pathhar chatta | *Garra gotyla* | Managers of Aquaculture-based Fish Hatchery at DVC, Jharkhand |
| 14. | Singhi | *Heteropneustes fossilis* | All |
| 15. | Silver carp | *Hypophthalmichthys molitrix* | All |
| 16. | Bighead, Briket, Brigate | *Hypophthalmichthys nobilis* | All |
| 17. | Bata | *Labeo bata* | All |
| 18. | Kalbas, Kalbos | *Labeo calbasu* | Staff of the Fisheries Department, Jharkhand<br>Members of the Fishermen Co-operative Society, Jharkhand |
| 19. | Rohu, Rui | *Labeo rohita* | All |
| 20. | Baami, Baam | *Mastacembelus armatus* | Staff of the Fisheries Department, Bihar<br>Members of the Fishermen Co-operative Society, Bihar |
| 21. | Tengra, Tangra | *Mystus tengara* | All |
| 22. | Patra, Falat, Pholui | *Notopterus notopterus* | All |
| 23. | Tilapia, Telapia, Jalebi | *Oreochromis* spp. | All |
| 24. | Pangaas, Bachua, Basa | *Pangasius pangasius* | All |
| 25. | Roopchand | *Piaractus brachypomus* | All |
| 26. | Punthi, Puthi | *Puntius* spp. | All |
| 27. | Ritha, Rita | *Rita rita* | Staff of the Fisheries Department, Bihar<br>Members of the Fishermen Co-operative Society, Bihar |
| 28. | Gagari | *Sperata seenghala* | All |
| 29. | Hilsa | *Tenualosa ilisha* | All |
| 30. | Mahseer | *Tor* and *Neolissochilus* spp. | Two Staff Members of the Fisheries Department, Jharkhand<br>One Researcher from Jharkhand |
| 31. | Boal, Bual, Buari | *Wallago attu* | All |

Only a small number of respondents from both states were aware of mahseers as freshwater fish in general, and even among these, perceptions regarding their presence varied considerably. In Jharkhand, respondents believed that the mahseer was either absent (4), endangered or possibly extinct in the rivers of the state (2; Fig. 3). Meanwhile, respondents in Bihar predominantly perceived mahseers to be absent (4) or present only in the cold-water rivers of the neighbouring country of Nepal (4; Fig. 4). Furthermore, the staff of the state fisheries (J: 9; B: 5) and forest departments (J: 3), and the Damodar Valley Corporation (DVC) fish hatchery (J: 2) reported that they did not possess any official records regarding the landing of the mahseers, though the possibility of occasional unofficial catch going unrecorded and unreported due to the protected status of these fish cannot be ruled out (Fisheries Dept. Jharkhand: 1).

As articulated in the following statement by a staff of the forest department from Jharkhand, the lack of systematic documentation of aquatic biodiversity and its distribution in the state (2) acts as a barrier for designing effective programmes for the conservation of native fish species.

> Quote 8: “Even though Jharkhand has so many jungles, rivers and wild animals, very little wildlife biodiversity research has been conducted here as compared to the other states of the country. On top of that, there has been even less research focused on aquatic biodiversity distribution or conservation. It is very important to undertake such kinds of studies to know the fish’s distribution. We are also willing to support researchers in doing so.”

These findings point towards an on-going process of “societal extinction” (Jarić et al. 2022), wherein species vanish not only from ecosystems but also from cultural memory. Because people no longer recognise or remember mahseer, there is little motivation to support its conservation (J: 64; B: 82). This result serves as a warning to the managers of aquatic ecosystems and conservators of biodiversity in Jharkhand and Bihar and other Indian states. If proper actions are not taken immediately other important fish species also may join mahseer in the journey towards “societal extinction” soon, since they seldom attract the notice of the public or policy makers due to the cryptic nature of their habitat and reduced opportunity for interaction. Therefore, strengthening the knowledge of diverse stakeholders and socio-cultural memory is essential to reinforce public interest, governmental support, policy attention and long-term conservation of native freshwater fishes including mahseers which may require restocking or reintroduction in Jharkhand and Bihar.

## 4. Discussion

Jharkhand and Bihar are among India‘s economically poorer states, where riverine and inland fisheries remain central to livelihoods, food security, and nutrition for millions (Dey et al. 2020; Kumar, V. et al. 2025). Hence, conservation in these regions is not driven solely by ecological decline, environmental challenges, or data gaps, but by the interlinked and deeply embedded socio-economic, institutional and behavioural dynamics that shape stakeholder priorities, decision-making and actions. Bihar has emerged as one of the leading inland fish-producing states with the largest population of inland fishermen in India, through the rapid expansion of pond aquaculture and hatchery-based seed production (Kumar et al. 2018; Rupala, Parliament of India 2023; Prasad 2025), while Jharkhand has promoted reservoir fisheries and cage culture, including large-scale installations such as those in Chandil reservoir in the Seraikela-Kharsawan district (Pandit et al. 2021; Kumari and Sharma 2022). Although, these interventions have enhanced fish yields, improved food security, and generated employment opportunities, it also reinforced instrumental values towards freshwater ecosystems and shifted management priorities towards production-oriented goals rather than conservation-centred actions (Kumari and Sharma 2022; Raut et al. 2020; Arora et al. 2024). As a result, long-term ecological sustainability and native fish biodiversity are frequently marginalised within governance frameworks, receiving limited conservation attention compared to aquaculture-driven development (Béné 2003; Deb 2009; Leite and Gasalla 2013; Lynch et al. 2016; Dey et al. 2020; Arora et al. 2024).

This production-centric approach, coupled with the expansion of aquaculture through the seed-production of non-native fishes, has led to significant ecological consequences. It has contributed to the widespread introduction and establishment of exotic IAFs, including tilapia (*Oreochromis* spp.), African catfish, common carp (*Cyprinus carpio*), grass carp (*Ctenopharyngodon idella*), silver carp (*Hypophthalmichthys molitrix*), and bighead carp (*Hypophthalmichthys nobilis*) across the freshwater-bodies in Jharkhand and Bihar (Canonico et al. 2005; Singh and Lakra 2006). These species often outcompete native fishes for resources, alter food chains and webs, and reduce ecosystem resilience (Khan and Panikkar 2009; Martin et al. 2010). Despite widespread declines in populations of native fish species, aquaculture-driven development has emerged as the dominant policy (Narayanan 2016; Kumari and Sharma 2022) over the sustainable management of natural fish populations (Arora et al. 2024; Kumar, V. et al. 2025) leaving conservation interventions for native fishes disjointed, less prioritised, with limited emphasis on ecosystem restoration, stakeholder participation, or long-term sustainability.

The prioritisation of livelihoods and economic benefits expressed by the fishermen, hatchery managers, and fisheries department staff does not reflect that they are against conservation per se, but rather highlights a mismatch between conservation goals and livelihood realities. Although stewardship values, positive attitude towards conservation and willingness (conditional) were present their conservation intentions remained weak because of the low perceived behavioural control (PBC). Stakeholders questioned the feasibility of investing time and resources in conservation due to the absence of social support or compensation for the time spent in such activities which may jeopardise their income. Such livelihood-first norms are not unique to Jharkhand and Bihar; similar patterns have been documented across many economically marginal regions, where conservation is often perceived as a luxury rather than a necessity (Béné 2003; Deb 2009; Haque 2009; Kiwango and Mabale 2022; He and Jiao 2023; Lyakurwa et al. 2025). Evidence from Asia (India, Bangladesh, rural China), Sub-Saharan Africa (Cameroon, Congo, Nigeria, Chad), and South America (Brazil) have consistently shown that livelihood insecurity compels local communities to prioritise short-term subsistence and income security over long-term ecological and resource sustainability. In such contexts, conservation interventions are often viewed as externally imposed rather than locally relevant and a threat to immediate food access and livelihoods, thereby leading to low compliance and limited long-term support (Béné 2003; Deb 2009; Haque 2009; Kiwango and Mabale 2022; He and Jiao 2023; Lyakurwa et al. 2025).

Despite the popularity of the relational and intrinsic values, as well as a sense of responsibility for protecting fishes and freshwater ecosystems in both states, local fishermen expressed willingness to support conservation activities provided their livelihood needs are met, benefits are equitably distributed and they are included in the decision-making processes. The coexistence of positive conservation values with the prioritisation of the fish based livelihood underscores a classic attitude-behaviour gap (Barr 2006; Muller et al. 2025). While attitudes and values are broadly supportive of conservation, actual behaviour is constrained by limited resources, weak institutional support, non-inclusion of inputs and no autonomy in decision making, social norms prioritising income, and governance failures (TPB components), thereby leading to limited pro-conservation behaviour (Ullah et al. 2021; Karimi and Ataei 2022). These observations highlight the need for conservation approaches that align ecological goals with livelihood incentives, rather than framing them as competing priorities (Pimid et al. 2022; Newing et al. 2024), especially in economically poor but biodiversity-rich regions (Salafsky and Wollenberg 2000; Nepal and Spiteri 2011; Mandoloma et al. 2025). Failing to do so risks making the subjective norms that support short-term economic viability over the coexistence and conservation of indigenous and native (fish) species the mainstream narrative in the society, thereby making future changes increasingly difficult (Hutton et al. 2005; Bennett et al. 2017).

Although the study areas are blessed with numerous native fishes, the majority of the stakeholders we contacted were familiar largely with the species of aquaculture and fisheries importance. This observed absence of native fish diversity in respondents‘ knowledge base may lead to gradual erosion of the ecologically important but economically less significant fish species from the public discourse and their memory (Jarić et al. 2022; Soga and Gaston 2022), as observed in the case of mahseers. The presence of these charismatic megafishes - *T. putitora*, *T. tor*, *T. mosal* and *N. hexagonolepis* - has been reported from 15 of the 24 districts of Jharkhand and 12 of the 38 districts of Bihar. Historical records, including the British-era Gazetteers of Bihar and Bengal (1906-1926), the Government of Bihar Gazetteers, and several books and research articles dating back to the early 1900s, provide descriptions of mahseers from the focal states (O‘Malley 1906, 1910, 1924, 1926; Roy Choudhury 1957; Kumar 1970; Talwar and Jhingran 1991; Sugnnan 1995; Desai 2003; Das 2005; Sinha Ray 2016). Furthermore, drawing on extensive archival and repository-based research on hunting, wildlife and conservation in colonial Bengal (1850-1947), Sinha Ray (2016) reported that mahseers occurred in the Damodar and Subarnarekha rivers of Jharkhand (then part of Bengal) as early as the 1850s, highlighting the long historical distribution of these species in the eastern Indian river systems. However, poor species recognition by the stakeholders, meager representation of these fishes in conservation education, minimal breeding programmes, and negligible representation in the social and conventional media (Das and Binoy 2024, 2025a) point towards a potential “dual (societal and biological) extinction” of mahseers in both states. This argument is reinforced by the absence of mahseers in the catch records of the last decades, unavailability of their distributional references in the recent research, and nonexistence of research programmes focusing these fishes since the late 1990s (Tudu and Maji 2025; Raghavan et al. 2017; Dahanukar et al. 2018; Pinder et al. 2019; Das and Khan 2024; Sarmah et al. 2024). When human interactions with a species decline, it loses societal salience and gradually disappear not only from the ecosystem but also from the cultural (art, literature, digital records) as well as communicative (lived experiences, local names, folklores; Assmann 2011; Candia et al. 2019; Jarić et al. 2022) memories. Such societal extinction can further diminish the possibility of getting support from public and policy makers for conservation efforts, such as the ecological exploration to know the presence, or undertake restocking programmes for, red-fin or deep-bodied mahseer (*T. tor*), mosal mahseer (*T. mosal*), golden mahseer (*T. putitora*), and chocolate mahseer (*N. hexagonolepis*), once common in the waters of Bihar and Jharkhand. These results offer an important lesson and a warning for other regions in India, as well as other nations harbouring natural populations of mahseer - if timely conservation interventions are not implemented, a similar situation is not far away!

Our results point towards an urgent need for the targeted communication and collaborative interventions in Jharkhand and Bihar to ensure the conservation of freshwater fish diversity, and counter the on-going erosion of socio-cultural memory associated with mahseer. Differing from the terrestrial organisms, due to the cryptic nature of their habitat and the reduced opportunity for interaction available to a major section of the human population, extermination of many freshwater fish species could happen silently (Cambray and Bianco 1998; Reid et al. 2019; Vardakas et al. 2025). Hence, starting ecological surveys, in-depth distribution studies, conducting multi-stakeholder workshops and awareness programmes, and establishing collaborations are the need of the hour to save this important component of biodiversity. In this context, the societal extinction faced by the mahseers, along with a biological one, in these two states could be projected as a compelling example to convince the policymakers and mobilise the public to prioritise and participate in fish conservation programmes. Notably, past attempts at wild mahseer population recovery very significantly highlight the consequences of inadequate planning. In the late 1990s, attempts were undertaken to reintroduce mahseer twice in the North Koel River of Jharkhand (then part of undivided Bihar). However, these efforts were not successful due to unsuitable ecological conditions, in terms of elevated river water temperatures, as well as the lack of adequate scientific guidance and planning (Singh *personal communication*). Similarly, during the year 2015-2016, the ICAR-DCFR, Bhimtal, supplied 1,000 *T. putitora* fingerlings to the Department of Fisheries, Bihar (ICAR-DCFR 2016). However, no further information or documents are available regarding the subsequent status of these fingerlings, including whether they were released into the natural freshwater water-bodies for stock enhancement. This absence of follow-up data, as well as the lack of further efforts to reintroduce the fingerlings, is not merely an administrative gap; it reflects a broader systemic issue concerning the prioritisation of fish conservation. Therefore, scientifically-backed reintroduction programmes, coupled with effective follow-up and long-term monitoring, alongside the promotion and dissemination of historical and cultural narratives, could help revive public interest and collective memory of mahseer (Jarić et al. 2022), and potentially extend the enthusiasm to other native fish species. Furthermore, the presence of an active communication system will preserve local knowledge, promote community involvement, improve compliance of conservation norms and laws (Ostrom 2009; Bennett et al. 2017), and reduce human-human conflict (Huang et al. 2020; Wu et al. 2020). Although The Bihar Fisheries Bill, 2013 (draft) was envisaged as a comprehensive legislative framework to promote conservation of aquatic biodiversity, sustainable utilisation of fish stocks, establishment of advisory committees for recommendations, declaration of fish sanctuaries, prohibition of destructive fishing gears and practices, scientific development of fisheries, streamlining licensing mechanisms, and improving the quality fish seed (Narayanan 2016; Kumar et al. 2019), thereby addressing the gaps existing in the Bihar Fish Jalkar Management Act, 2006 (Bihar Act, 13 of 2006, amended in 2018, thereby called The Bihar Fish Jalkar Management (Amendment) Act, 2018), it remained at the draft stage. The enactment of this bill as a stand-alone Act (Kumar et al. 2019), along with the development of similar fisheries legislative acts and policy frameworks for Jharkhand, was also recommended by several stakeholders to reduce the loss of fish diversity.

Targets 21 and 22 (Section H) and Section K of the Kunming-Montreal Global Biodiversity Framework (CBD 2022), emphasise participatory and inclusive decision-making, and recognise the critical role of communication strategies that extend beyond just awareness-raising to foster positive and sustained behavioural change towards the biodiversity (CBD 2022). Unfortunately, across both states, governance structures were characterised by top-down decision-making, minimal stakeholder participation and limited communication and collaboration among government departments, researchers and local fishermen communities. The *Matsya Mitra* (‗Friends of Fishes‘) initiative undertaken by the Jharkhand State Government in 2007, offer a distinct example of how community-based intermediaries can improve communication, trust, and knowledge transfer (Shweta 2024). The *Matsya Mitra* are people from the local communities appointed on a voluntary basis to work as bridges and communicators between the fisheries department and local community fishermen (NITI Aayog n.d.; Shweta 2024). Although, their primary focus is the promotion of aquaculture, including conservation education and creating awareness of the native fish diversity in their mandates, and empowering them to lead the monitoring and restoration of the freshwater habitats, could help to enhance stakeholder ownership and trust amongst the local community members (Schultz and McGinn 2013).

Taken together, freshwater fish conservation in Jharkhand and Bihar appeared to be constrained less by the dearth of positive attitude and more by governance and communication gaps, limited stakeholder and community participation, misaligned incentives, and critical weaknesses in the decision-making systems, as revealed during the decision-making and implementation stages of CPF. From a TPB perspective, these institutional barriers significantly curb the perceived behavioural control, even among stakeholders keen to participate in conservation activities (Zolait 2014; Dong et al. 2024). Active fishermen‘s cooperative societies in Jharkhand curbing illegal destructive fishing practices suggests that collective action and involvement of local groups can translate relational tribal values (Jolly et al. 2022; Iqbal et al. 2025; Jolly and Stronza 2025; Kumar, A. et al. 2025) into conservation positive behaviour and endorse human-animal coexistence if proper supports are provided. Our observations resonate with emerging literature emphasising the role of relational values in conservation, where people‘s sense of responsibility, care, and connection to nature and wildlife motivates stewardship when structural barriers are addressed (Tsing 2012; Ingold 2013; Chan et al. 2016; Haraway 2018; Pascual et al. 2023; Murali et al. 2024; Jolly and Stronza 2025). Hence, strengthening the co-management of freshwater ecosystems, and recognising local ecological knowledge along with the lived experiences of tribal and local community members, can help translate the strong pro-conservation attitudes present among the stakeholders into sustained pro-environmental behaviours (Kelkar and Arthur 2022; Kura et al. 2023; Newing et al. 2024; Das and Binoy 2025b). However, unequal distribution of the benefits from the government *yojanas* (schemes) reported from the Bihar, if not managed effectively, could not only erode the legitimacy and acceptance of the policies (Bavinck et al. 2018; Owusu and Adjei 2021) but will also intensify the caste and politics based negotiations for traditional fishing rights and other forms of human-human conflict (Bagchi 2018; Kelkar and Arthur 2022), thereby weakening fish conservation efforts.

### Stakeholder Recommendations

In Jharkhand and Bihar, stakeholders‘ recommendations focused on practical, on-ground actions. These included organising regular meetings and encouraging discussions between fishermen and the fisheries department staff to address the challenges faced at the local level, and undertaking more scientific studies to document aquatic biodiversity distribution and support conservation. The declaration of fish sanctuaries and no-fishing zones, the promotion of conservation education and awareness programmes with a focus on native fish species, and the inclusion of inputs from the fisheries departments into research initiatives undertaken by the central government were also emphasised by many. Incorporating mahseer ecology and the socio-cultural significance of these species into the curricula of educational institutions focused on fisheries and aquatic sciences, as well as the role of such institutions in collecting and preserving voucher specimens of important native species also emerged during the discussion. Additional measures for protecting native fish populations proposed by numerous stakeholders included reviving breeding grounds and restoring habitats for native fish species, removing obstructions such as *bunds* (or *bandhs*) from the rivers, and ensuring that the benefits of government schemes reach all fishermen. These actions were seen by the respondents as essential for strengthening livelihood opportunities and fostering long-term local support for the conservation efforts.

## 5. Conclusion

While many stakeholders expressed positive attitudes towards conserving native fishes in the resource-limited Indian states of Jharkhand and Bihar, dominant subjective norms prioritising livelihood security over conservation combined with low perceived behavioural control curtailed its translation into meaningful conservation actions. Our findings underscore the need to move beyond purely ecological and taxonomic approaches, prominent in the conservation research, toward planning grounded in the socio-cultural uniqueness of the focal species or ecosystems. In both the focal states, freshwater fish conservation must be embedded within livelihood systems, inclusive institutional and decision-making structures, and local value frameworks. The integrated CPF-TPB-SV framework revealed that enabling conservation of the freshwater fishes will require interventions across multiple levels: (i) reshaping subjective norms by linking conservation with livelihoods, (ii) enhancing perceived behavioural control through incentives (monetary or non-monetary in terms of daily grocery and kitchen-related items or symbolic recognition (public visibility, media coverage); Kipkeu et al. 2014; Rode et al. 2015; Ewane 2024), capacity-building, and inclusive governance and decision-making, (iii) leveraging existing stewardship and relational values through bottom-up partnerships, acknowledgement and resolution of local needs and concerns, and transparent and equitable benefit-sharing, (iv) promoting collaborations between locals and researchers, and (v) rebuilding societal memory (cultural and communicative) of the mahseer, through targeted distribution research, historically-linked awareness and education. Any efforts to protect the native fish diversity in Jharkhand and Bihar that fail to address these socio-cultural and economic dimensions are less likely to succeed and risk remaining just symbolic, fragmented and ineffective, regardless of ecological urgency.

## Author Contributions

**Prantik Das:** Conceptualisation; methodology; investigation; data curation; formal analysis; visualisation; funding acquisition; writing - original draft; writing - review and editing. **V. V. Binoy:** Conceptualisation; methodology; visualisation; writing - review and editing; supervision.

## Acknowledgments

Prantik acknowledges the Human Research Development Group (HRDG) - Council of Scientific and Industrial Research (CSIR), New Delhi, India (09/1320(0001)/2020-EMR-I) for providing the research fellowship and the annual contingency grant for field-work related expenses. He extends his sincere appreciation and gratitude to all the respondents who took part in the interviews across districts in both study states. He also expresses his gratitude to Mrs. Bubu Sarkar Das and Mr. Subhashis Das for their assistance in facilitating stakeholder contacts and data collection in Jharkhand, and to Mr. Shivaditya Aich, Mr. Asad Bilal and Ms. Anjali Bharati in Bihar.

## Ethical Statement

All interviews were undertaken in compliance with the institutional ethical guidelines for conducting non-invasive research involving human participants, as approved by the Institutional Ethics Committees (IEC) of The University of Trans-Disciplinary Health Sciences and Technology (TDU; Protocol Number: TDU/IEC/16E/2025/PR72) and the National Institute of Advanced Studies (NIAS; Letter Number: NIAS-EC-11/03/2022).

## Informed Consent Statement

Prior to the participation, all respondents were made fully aware about the objectives of the study, and it was emphasised that their participation was completely voluntary, with the option to withdraw from interviews at any stage. The data collection through audio-recordings, note-taking and photography was carried out only after obtaining oral/audio-recorded informed consent. The participants were also assured that their names and other personal details that could be used to identify them, would remain confidential and anonymous to protect their identity and privacy.

## Data Availability Statement

In order to maintain the respondent‘s privacy, the datasets have not been made publicly available, as they contain information that could compromise research participant consent. Accordingly, the data used in this study remain confidential.

## Supplementary Materials (SM)

SM 1. Stakeholder Interview Questions:

*Section 1: Conservation Planning Framework (CPF)*

1. Are you aware of the different fish species found in your state?
2. What threats do you think that the freshwater fishes face in your locality?
3. Do people practice any destructive fishing methods here? What are those?
4. Do you know what the Invasive Alien Fishes (IAFs) are? Do you think that IAFs have any effect on the native fish populations in your locality?
5. Should there be stocking of IAF in rivers where native fish population thrives for the economic benefit? Why/Why not?
6. Do you think that there is a need to conserve the native fish species? If so, any opinions on which species need more attention?
7. What are the existing freshwater native fish population management programmes that are being undertaken by the Department (Fisheries and Forest) in the state?
8. Do you think these have been effective in leading to a positive outcome? Do you feel any need for change in these programmes?
9. Do you think interventions are required for the conservation of the native fish populations? What interventions can be done to bolster the stock in the wild?
10. Is subsistence fishing allowed for the locals within the National Park/Wildlife Sanctuary/protected area?
11. Have the local community members or other stakeholders (such as researchers, fishermen) been involved in the decision making process related to native fish conservation plans and strategies?
12. Does the Department (Fisheries and Forest) hold open discussions regarding fish conservation with the local community members?
13. Does the stakeholder hold discussions with the local community members or involve them in awareness activities?
14. Does the Department (Fisheries and Forest) work in association with other government Departments with respect to fish conservation programmes?
15. Does the stakeholder work in association with the other stakeholders such as research institutions, NGOs?
16. Are there sufficient resources (human and monetary) for dedicated fish conservation?
17. What are the conservation communication and awareness activities being undertaken by the stakeholders? Do the locals support these activities?
18. How important do you think conservation communication is for native fish conservation activities to be successful?
19. Does the local media focus on the conservation of native fish species?
20. Do you think social media can be used as a tool for communication?
21. What is your opinion on the creation of fish sanctuaries or no fishing green zones?
22. Are you aware of the mahseer fish species found in your state? What are they called in the local language?
23. (If unfamiliar with the term ‗mahseer,‘ photographs of different mahseer species were shown). Do you recognise these fishes?
24. (If unaware) Based on historical records, they were present in this region in the past. Is it correct that they occurred here earlier but are not found currently?
25. Do you think the number of mahseer (species name changed according to the state where the interview is being conducted) is declining in the rivers in your locality?
26. Are you aware about the IUCN status of the mahseer species?

*Section 2: Theory of Planned Behaviour (TPB)*

1. In your view, do the stakeholders possess adequate knowledge and information to effectively undertake fish conservation efforts on their own? Is that knowledge shared with other stakeholders?
2. Is there any kind of social pressure from your colleagues/government/family members/head of the community to involve in native fish conservation or other activities regarding fishes?
3. In your opinion, does the existing official workload limit the capacity of the staff to focus specifically on native fish conservation efforts? (For Government stakeholders)
4. What do native fishes mean to you, do you either see it from the point of view of conservation or a way for economic benefits or a combination of both?
5. Are you involved with conservation activities? If not, how willing would you be to get involved in conservation initiatives in your area?
6. Do you collaborate with other stakeholders in such activities?
7. What personal needs, interests, or motivations might drive you to get involved in these kinds of initiatives?
8. What would you say, in your opinion, are the barriers to native fish conservation?

*Section 3: Social Values (SV)*

1. Are the local native fishes a part of any mythological stories?
2. Do fishes in your locality have any socio-cultural or religious significance?
3. (If yes) In your view, should such cultural values be integrated into the design of region-specific fish conservation plans, such as temple fish sanctuaries? How effective would those be?
4. What kind of socio-economic benefits do native fishes provide to the community over generations?
5. Why do you think that the native fishes should be protected and conserved?
6. What kind of non-tangible, non-monetary benefits, in your opinion, do these native fishes provide to you and the society?
7. How do you think stakeholders view fish conservation — as a personal commitment to nature and biodiversity conservation, a livelihood opportunity, a reflection of cultural or religious identity, a job responsibility (professional obligation), or a combination of these?
8. Do you think that the stakeholders should feel that they have a duty towards the fish conservation?
9. Are there any additional recommendations you would like to suggest to enhance the effectiveness and efficiency of the existing native fish conservation protocols and strategies?

***Additional questions for Fisheries Department staff:***

1. (For Bihar) Are there any provisions for the declaration of Fish Sanctuaries in accordance with the Bihar Jalkar Management Act, 2006 or any other fisheries Acts or rules? Have they been declared?
2. If yes, what are those stretches? Who manages them?
3. What kind of human activities are allowed here?
4. (For Jharkhand) Have there been any provisions to declare river stretches as Fish Sanctuaries?
5. What are the activities done in the hatchery? (breeding, rearing, selling, distribution without cost)
6. Which species are being bred in the hatchery?
7. Are mahseers being bred in the hatchery? Have they been bred in the past? Any future plans to breed them?
8. Do the officials train the fish farmers regarding fish conservation or only for IMC aquaculture?
9. Do you provide fingerlings to local people? For what purpose?
10. Is there fish conservation related breeding that is being undertaken? (If no) Why not?
11. (For Jharkhand) What is the role of *Matsya Mitra*?

***Additional questions for Forest Department staff:***

1. Are there any targeted native fish conservation plans or are they protected as part of the forest and biodiversity?
2. Do you conduct a creel census to determine the number of individuals?
3. Are the forest guards consulted during discussions with the ranger or DFOs?
4. Do the forest guards face any kind of conflict from the local residents of the buffer zone?
5. Are the local community members invited for discussions over decision making?
6. Are the buffer zone residents permitted to subsistence fish with rod without any destructive methods? Were the rules the same before or have they changed?
7. Do you provide any licenses to local fishermen for fishing in the buffer zone of the protected areas? If yes, how many licenses are provided? Who are they provided to?
8. Is there any closed season for fishing?
9. What are punishments for illegal destructive fishing or poaching?

***Additional questions for researchers:***

1. Are you invited to discussions over native fish conservation and management decision making?
2. Are you involved in native fish conservation research?
3. Does your research have a component of intervention as well?
4. Have you conducted any intervention? How effective have those been?
5. Are there any barriers in breeding native fish species?
6. Have you been consulted during breeding or reintroduction or stocking programmes for conservation purposes?
7. Have you faced any conflict with any stakeholder during your research work which included field-work?

***Additional questions for fishermen:***

1. What kind of fishes do you generally catch?
2. What kind of fishes do you farm for aquaculture?
3. Are you entirely dependent on fishing and aquaculture for your livelihood?
4. Do you need a license to fish in certain stretches of the river? What are these stretches? Who issues the license?
5. Have the number of native fish caught declined over the years? Do you get IAFs in your catch? Have they increased?
6. Do you catch mahseer? If yes, do you land them frequently?
7. What do you do if you land a mahseer? Are they taken for consumption purposes?
8. Are you consulted by the fisheries department officials on what kind of fish to be bred?
9. Do discussions happen with the fisheries department?
10. Are you involved in any native fish conservation related activities?
11. How important is involvement in conservation activities for you along with the livelihood based fishing and aquaculture?
12. Have you seen destructive illegal fishing being undertaken? Have they declined or increased over time?
13. Along with training regarding IMC breeding and aquaculture, are you provided with training or education regarding conservation of native fish species?
14. In case you are asked to involve yourself in conservation activities, will you be willing to do so? Are there sufficient people for undertaking such activities?
15. Do you receive regular benefits from the government-based schemes (*yojanas*) that are in place for the welfare of the fishermen community?
16. Do you face any barriers in receiving them?
17. In your opinion, how can the native fish conservation and livelihood-based aquaculture activities go hand-in-hand and promote sustainability?

